# Lipid availability influences the metabolic maturation of human pluripotent stem cell-derived cardiomyocytes

**DOI:** 10.1101/2020.03.14.991927

**Authors:** Hui Zhang, Mehmet G. Badur, Sean Spiering, Ajit Divakaruni, Noah E. Meurs, Michael S. Yu, Alexandre R. Colas, Anne N. Murphy, Mark Mercola, Christian M. Metallo

## Abstract

**Objectives:** Pluripotent stem cell-derived cardiomyocytes are phenotypically immature, which limits their utility in downstream applications. Metabolism is dramatically reprogramed during cardiac maturation *in vivo* and presents a potential avenue to drive *in vitro* maturation. We aimed to identify and address metabolic bottlenecks in the generation of human pluripotent stem cell (hPSC)-derived cardiomyocytes.

**Methods:** hPSCs were differentiated into cardiomyocytes using an established, chemically-defined differentiation protocol. We applied 13C metabolic flux analysis (MFA) and targeted transcriptomics to characterize cardiomyocyte metabolism in during differentiation in the presence or absence of exogenous lipids.

**Results:** hPSC-derived cardiomyocytes induced some cardiometabolic pathways (i.e. ketone body and branched-chain amino acid oxidation) but failed to effectively activate fatty acid oxidation. MFA studies indicated that lipid availability in cultures became limited during differentiation, suggesting potential issues with nutrient availability. Exogenous supplementation of lipids improved cardiomyocyte morphology, mitochondrial function, and promoted increased fatty acid oxidation in hPSC-derivatives.

**Conclusion:** hPSC-derived cardiomyocytes are dependent upon exogenous sources of lipids for metabolic maturation. Proper supplementation removes a potential roadblock in the generation of metabolically mature cardiomyocytes. These studies further highlight the importance of considering and exploiting metabolic phenotypes in the *in vitro* production and utilization of functional hPSC-derivatives.

## 1. Introduction

The regenerative capacity of human pluripotent stem cells (hPSCs) can provide a virtually unlimited source of cardiomyocytes (CMs) and other functional cells for toxicology and/or tissue engineering applications (Ye et al., 2014, Chong et al., 2014, Takeda et al., 2018, Matsa et al., 2014). Recently developed cardiac differentiation protocols now generate CM purities approaching >90% without the use of growth factors in partially- or fully-defined conditions (Lian et al., 2012, Burridge et al., 2014). However, differentiated CMs can be immature such that they lack the proper electrical connectivity, force generation, and metabolism to properly mimic their *in vivo* counterparts (Yang et al., 2014, Kolanowski et al., 2017). Preclinical models of CM transplantation have shown potential increased arrhythmia risk and demonstrated the need for functional maturation (Liao et al., 2010, Zhang et al., 2002). In particular, recapitulation of native mitochondrial function and bioenergetics will be essential to effectively make use of hPSC-derived CMs.

While aging of CMs has generated some success (Lundy et al., 2013), this is impractical for bioprocess scale up and quantities needed eventually *in vivo* after myocardial infarction (i.e. delivery of up to 10^9^ cells) (Behfar et al., 2014, Chong et al., 2014). Efforts to speed up functional maturation of CMs *in vitro* have focused on mechanical cues (Ronaldson-Bouchard et al., 2018), electrical stimulation (Radisic et al., 2004, Nunes et al., 2013), and physical microenvironment (Ribeiro et al., 2015, Shadrin et al., 2017, Ulmer et al., 2018); however, manipulation of the nutrient environment/culture medium provides a promising avenue for promoting metabolic maturation of hPSC-CMs (Tohyama et al., 2013).

The bioenergetic demands of the developing and adult heart require a dramatic upregulation of ATP regeneration, which is a marked by a transition from highly glycolytic progenitors to functional CMs that ultimately rely on oxidative phosphorylation for energy generation (Gaspar et al., 2014). Recent work has demonstrated that this mitochondrial shift is necessary for proper CM development (Hom et al., 2011). Moreover, the increase in mitochondrial activity is marked by the catabolism of diverse carbon sources in the adult heart that drives proper CM function in the fasted and fed states (e.g. fatty acids; ketone bodies; branched-chain amino acids, BCAAs) (Lloyd et al., 2004, Kolwicz et al., 2013). As with other measures of CM function, the metabolism of hPSC-derived CMs is fetal-like, with heavy reliance on glycolysis and glucose oxidation, and presents a roadblock for their utility (Lopaschuk and Spafford, 1990, Razeghi et al., 2001). However recent work has definitively demonstrated that activating mitochondrial function through galactose supplementation induces a more mature CM metabolism (i.e. fatty acid oxidation) and better function in hPSC-derived CMs (Correia et al., 2017). The role of mitochondrial substrate switching during CM differentiation and the role of fatty acid metabolism during maturation is underexplored.

Here we have examined hPSC-CM metabolism throughout differentiation to identify key pathways activated to fuel CM bioenergetics. Immature hPSC-CMs activate pathways such as ketone body and BCAA oxidation while suppressing glutaminolysis. However, these cells fail to activate fatty acid oxidation (FAO) during differentiation. Day-by-day tracing revealed an activation of enzyme expression and mitochondrial catabolism of key substrates during differentiation, suggesting a correct differentiation program and some other obstacle to proper CM function. Further analysis revealed aberrant expression of genes involved in lipid oxidation and biosynthesis, leading us to hypothesize that lack of fatty acid supplementation in differentiation media forces CMs to synthesize structural lipids rather than oxidizing them for fuel. Supplementation of complex fatty acid mixtures improved mitochondrial function and substrate oxidation without compromising CM electrical function. Together these data highlight the importance of hPSC-nutritional microenvironments during maintenance and differentiation to facilitate normal physiologic functions of their derivatives.

## 2. Materials and Methods

### 2.1. Human pluripotent stem cell (hPSC) culture

Human embryonic stem cell line WA09 (H9) and induced pluripotent stem cell line iPS(IMR90iPS)-c4 were supplied by WiCell Research Institute. HPSCs were routinely maintained in mTeSR1 media (Stem Cell Technologies) on growth factor–reduced Matrigel (Corning Life Sciences) at 8.8*µ*g/cm^2^ and passaged every 4 days using ReLeSR (Stem Cell Technologies). All hPSCs experiments were conducted with cells ranging from 40 and 70 passages. For metabolic tracing experiment, hPSCs were adapted to chemically defined media TeSR-*E8* media (Stem Cell Technologies) for at least one passage, and all hPSCs were detached and plated by exposure to Accutase (Innovative Cell Technology).

### 2.2. Cardiomyocyte (CM) differentiation

All hPSCs were cultured for at least five passages post thaw before beginning differentiation. HPSCs were differentiated by the adapted chemically defined CM generation protocol (Burridge et al., 2014). Briefly, hPSCs were dissociated using a 0.5mM EDTA (Life Technologies) in PBS without CaCl_2_ or MgCl_2_ (Corning Life Sciences) for 7 minutes at room temperature. HPSCs were plated at about 3.0×10^5^ cells/cm in mTeSR1 or TeSR-*E8* media (Stem Cell Technologies) supplemented with 2 μM Thiazovivin (Selleck Chemicals) for the first 24 hours after passage. HPSCs were fed for 3-5 days until they reached ≥ 90% confluence. To initiate differentiation, cells were washed with PBS 1X and the culture medium was changed to CDM3, consisting of RPMI 1640 medium (Life Technologies), 500μg/ml *O. sativa*–derived recombinant human albumin (Sigma-Aldrich), and 213μg/ml L-ascorbic acid 2-phosphate (Sigma-Aldrich), with 6 μM CHIR 99021 (Selleck Chemicals) for 48 hours. Media was then changed to CDM3 with 2 μM Wnt Inhibitor C59 (Selleck Chemicals) for 48 hours. After 48 hours this medium was replaced with basic CDM3 media. 48 hours later cells were then dissociated with TrypLE Express (Life Technologies) and plated onto Matrigel coated plates in CDM3 supplemented with 200nM Thiazovivin at a density of 8.9×10^5^ cells/cm^2^ to generate well to well homogeneity 48 hours after replating, media was changed to either basal CDM3 media or CDM3 + Albumax/ 0.5nM L-carnitine media (name?). The media was completely replaced every 48 hours thereafter. Spontaneous beating was typically observed8 to 10 days after differentiation was initiated. On differentiation day 16 media was replaced with Glucose free CDM3 medium with 10 mM sodium DL-lactate (Sigma-Aldrich), was used for further *in vitro* CM purification (2 to 4 days). Cells were then maintained in CDM3 or CDM3+Albumax+L-carnitine with media changes every 48 hours.

### 2.3. 13C metabolic tracing

For tracer experiments, culture medium was removed, cells were rinsed with PBS, and tracer media was added to wells. Tracer media consisted of glutamine, glucose, amino acid-free RPMI1640 (US Biologics) supplemented with proper levels of ^12^C amino acids, organic acids, and carbohydrates not being traced. If a more replete condition was being tested (i.e. additional nutrients added to basal media), ^12^C metabolites were added to other tracer arms to ensure equivalent nutrient state (i.e. add ^12^C lactate to a ^13^C glucose trace). The following tracers were used: [U-^13^C_5_]glutamine (Cambridge Isotope Laboratory), [U-^13^C_6_]glucose (Cambridge Isotope Laboratory), [3-^13^C]pyruvate (2mM; Cambridge Isotope Laboratory), [3-^13^C]lactate (2mM; Cambridge Isotope Laboratory), [U-^13^C_6_]leucine (Cambridge Isotope Laboratory), [U-^13^C_4_]β-hydroxybutyrate (2mM; Sigma-Aldrich), [1-^13^C]octanoate (Sigma-Aldrich), or [U-^13^C_16_]palmitate (Cambridge Isotope Laboratory). [U-^13^C]palmitate was first non-covalently conjugated to ultra fatty acid free BSA (Roche) as previously described (Vacanti et al., 2014). BSA conjugated [U-^13^C_16_]palmitate was added to each culture medium along with additional L-carnitine at 50 *µ*M (5% v/v) and 0.5 mM, respectively.

### 2.4. Metabolite extraction and derivatization

Cellular metabolites and fatty acids were extracted using methanol/water/chloroform. Briefly, spent media was removed, and cells were rinsed with 0.9% (w/v) saline and 500 μL of −80°C MeOH was added to quench metabolism. 200 μL of ice-cold water containing 5 μg/mL norvaline (internal standard for organic acids and amino acids) was added to each well. Both solution and cells were collected via scraping. Cell lysates were transferred to fresh Eppendorf tube and 500 μL of −20°C CHCl_3_ containing 2 μg/mL D_31_-palmitic acid (internal standard for fatty acids) was added. After vortexing and centrifugation, the top aqueous layer and bottom organic layer were collected and dried.

Derivatization of aqueous metabolites was performed using the Gerstel MultiPurpose Sampler (MPS 2XL). Dried aqueous metabolites were dissolved in 20 *µ*L of 2% (w/v) methoxyamine hydrochloride (MP Biomedicals) in pyridine and held at 37°C for 60 minutes. Subsequent conversion to their tert-butyldimethylsilyl derivatives was accomplished by adding 30*µ*L *N*-methyl-*N*-(tert-butyldimethylsilyl) trifluoroacetamide + 1% tert-butyldimethylchlorosilane (Regis Technologies) and incubating at 37°C for 30 minutes. Fatty acid methyl esters (FAMEs) were generated by dissolving dried fatty acids in 0.5 mL 2% (v/v) methanolic sulfuric acid (Sigma-Aldrich) and incubating at 50°C for 2 hours. FAMEs were subsequently extracted in 1 mL hexane with 0.1 mL saturated NaCl. Cholesterol was derivatized from dried FAME fraction by first dissolving in 20*µ*L pyridine and then adding 30 *µ*L N-methyl-N-trimethylsilyl-trifluoroacetamide (Macherey-Nagel) at 37°C for 30 minutes.

### 2.5. GC/MS analysis

Gas chromatography/mass spectrometry (GC/MS) analysis was performed using an Agilent 7890A with a 30 m DB-35MS capillary column (Agilent Technologies) connected to an Agilent 5975C MS. GC/MS was operated under electron impact (EI) ionization at 70 eV. One microliter sample was injected in splitless mode at 270°C, using helium as the carrier gas at a flow rate of 1 ml/min. For analysis of organic and amino acid derivatives, the GC oven temperature was held at 100°C for 2 minutes, increased to 255°C at 3.5°C/min, then ramped to 320°C at 15°C/min for a total run time of approximately 50 minutes. For measurement of FAMEs, the GC oven temperature was held at 100°C for 3 minutes, then to 205°C at 25°C/min, further increased to 230°C at 5°C/min and ramped up to 300°C at 25°C/min for a total run time of approximately 15 minutes. For measurement of cholesterol, the GC oven temperature was held at 150°C for 1 minute, then to 260°C at 20°C/min, then held for 3 minutes, and further increased to 280°C at 10°C/min and held for 15 minutes, finally ramped up to 325°C for a total run time of approximately 30 minutes. The MS source and quadrupole were held at 230°C and 150°C, respectively, and the detector was operated in scanning mode, recording ion abundance in the range of 100 – 650 m/z.

### 2.6. Mole percent enrichment calculation

Mole percent enrichment (MPE) of isotopes was calculated as the percent of all atoms within the metabolite pool that are labeled: 

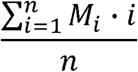

where *n* is the number of carbon atoms in the metabolite and M_*i*_ is the relative abundance of the *i*^th^ isotopomer. Citrate MPE derived from different nutrient tracers represent the flux of labeled nutrients into cellular central carbon metabolism.

### 2.7. Isotopomer spectral analysis (ISA)

For quantification of metabolites and mass isotopomer distributions, selected ion fragments were integrated and corrected for natural isotope abundance using MATLAB-based algorithm and metabolite fragments listed in Table 1. Total abundances were normalized by counts of internal standard control. Isotopomer spectral analysis (ISA) for quantitation of ^13^C contribution to lipogenic AcCoA was calculated as previously described (Vacanti et al., 2014). Specifically, the relative enrichment of the lipogenic AcCoA pools from a given tracer and the percentage of newly synthesized fatty acids were estimated from a best-fit model using the INCA MFA software package (Young, 2014). The 95% confidence intervals for both parameters were determined by evaluating the sensitivity of the sum of squared residuals between measured and simulated palmitate mass isotopomer distributions to small flux variations (Antoniewicz et al., 2006).

**Table 1.** Metabolite fragments used for GC/MS analysis.

| Metabolite | Abbreviation | Derivatization | m/z | Fragments for integration |
| --- | --- | --- | --- | --- |
| α-Ketoglutarate | aKG | tBDMS | 346 | $\text{C}_{14}\text{H}_{28}\text{O}_5\text{NSi}_2$ |
| Alanine | Ala | tBDMS | 260 | $\text{C}_{11}\text{H}_{26}\text{O}_2\text{NSi}_2$ |
| Aspartate | Asp | tBDMS | 418 | $\text{C}_{18}\text{H}_{40}\text{O}_4\text{NSi}_3$ |
| Lactate | Lac | tBDMS | 261 | $\text{C}_{11}\text{H}_{25}\text{O}_3\text{Si}_2$ |
| | | | 233 | $\text{C}_{10}\text{H}_{25}\text{O}_2\text{Si}_2$ |
| Citrate | Cit | tBDMS | 459 | $\text{C}_{20}\text{H}_{39}\text{O}_6\text{Si}_3$ |
| Fumarate | Fum | tBDMS | 287 | $\text{C}_{12}\text{H}_{23}\text{O}_4\text{Si}_2$ |
| Glutamate | Glu | tBDMS | 432 | $\text{C}_{19}\text{H}_{42}\text{O}_4\text{NSi}_3$ |
| Glycine | Gly | tBDMS | 246 | $\text{C}_{10}\text{H}_{24}\text{O}_2\text{NSi}_2$ |
| Malate | Mal | tBDMS | 419 | $\text{C}_{18}\text{H}_{39}\text{O}_5\text{Si}_3$ |
| Norvaline | | tBDMS | 288 | $\text{C}_{13}\text{H}_{30}\text{O}_2\text{NSi}_2$ |
| Proline | Pro | tBDMS | 330 | $\text{C}_{16}\text{H}_{36}\text{O}_2\text{NSi}_2$ |
| Pyruvate | Pyr | tBDMS | 174 | $\text{C}_6\text{H}_{12}\text{O}_3\text{NSi}$ |
| Serine | Ser | tBDMS | 390 | $\text{C}_{17}\text{H}_{40}\text{O}_3\text{NSi}_3$ |
| Succinate | Suc | tBDMS | 289 | $\text{C}_{12}\text{H}_{25}\text{O}_4\text{Si}_2$ |
| Oleate | Olea | FAME | 296 | $\text{C}_{19}\text{H}_{36}\text{O}_2$ |
| Palmitate | Palm | FAME | 270 | $\text{C}_{17}\text{H}_{34}\text{O}_2$ |
| Stearate | Stear | FAME | 298 | $\text{C}_{19}\text{H}_{38}\text{O}_2$ |
| Cholesterol | Chol | TMS | 458 | $\text{C}_{33}\text{H}_{54}\text{OSi}$ |

### 2.8. Oxygen Consumption Measurement

Respiration was measured in viable hPSC-derived CMs using a Seahorse XF96 Analyzer. HPSC-derived CMs were assayed in fresh culture media. ATP-linked respiration was calculated as the oxygen consumption rate sensitive to 2 mg/ml oligomycin in each culture condition and normalized by cell abundance. Each culture condition sample had at least four biological replicates analyzed. Cell abundance was indicated by the total fluorescence after staining with Hoechst 33342 (Divakaruni et al., 2014).

### 2.9. Gene expression analysis

Total mRNA was isolated from cells using MirVana kit for RNA extraction per the manufacturer’s protocol. Isolated RNA was reverse transcribed using Qiagen kit for cDNA synthesis per the manufacturer’s protocol. Real-time PCR (RT-PCR) was performed using SYBR green reagent (iTaq Universeal SYBR Green Supermix) per the manufacturer’s protocol. Relative expression was determined using Livak (ΔΔC_T_) method with *GAPDH* as housekeeping gene. Primers used are tabulated in supporting information and were taken from Harvard Primer bank (Wang et al., 2012).

### 2.10. Immunocytochemistry

All hPSCs were harvested and resuspended in 1% (v/v) paraformaldehyde and then fixed in 90% cold methanol. Cell pellets were incubated with a 1:200 dilution of human cTnT primary mouse antibody (ThermoFisher) overnight at 4°C. The solution was removed, cell pellets washed with PBS, and incubated with a 1:1000 dilution of the secondary antibody conjugated with Alexa Fluor® 488 at room temperature for 30 minutes. The cTnT+ cells were detected by BD flow cytometer. For microscopy adherent cells were fixed in 4% (v/v) paraformaldehyde. Cells were incubated with a 1:200 dilution of human cTnT primary mouse antibody for 1 hour at room temperature. The solution was removed, cells were incubated with a 1:1000 dilution of the secondary antibody conjugated with Alexa Fluor® 488 at room temperature for 20 minutes. Cells were subsequently washed and incubated with DAPI nucleus staining solution at room temperature for 15 minutes. Images at 20X were captured with a Zeiss fluorescent microscope.

### 2.11. Transmission electron microscopy (TEM) imaging

All hPSC-derived CMs were maintained on 12 well plates and collected on plates as clusters by scraping. Cell clusters were fixed with 2% paraformaldehyde and 2.5% glutaraldehyde in 0.15M sodium cacodylate buffer (SC buffer, pH 7.4). Samples were further incubated with 1% osmium in 0.15M sodium cacodylate for 1 to 2 hours on ice; washed 10 minutes in 0.15M SC buffer followed by rinsing in ddH_2_O on ice for 5 times; incubated in 2% of *uranyl acetate* for 1 to 2 hours at 4°C; dehydrated for 10 minutes in ETOH at 50%, 70%, 90%, 100% (twice) on ice; dried in acetone for 15min at room temperature; incubated in 50:50 (v/v) ETOH: Durcupan for at least one hour at room temperature; incubated in 100% Durcupan over night; after 24 hours, changed once in fresh 100% Durcupan for half day at room temperature; embed tissues in Durcupan in 60 °C oven for 36 to 48 hours. Ultrathin sections (60nm) were cut on Leica microtome with Diamond knife and followed by post staining with both uranyl acetate and lead. Images were captured at required magnification on FEI Spirit Tecnai TEM at 80KV with Eagle 4kx4k camera

### 2.12. Preparation of VF2.1.Cl loading solution and reference compounds

Physiological analysis was performed using a VF2.1.Cl dye, a newer version of Fluovolt and previously published method (McKeithan et al., 2017). Voltage dye loading solution and the compound dilutions were prepared prior to manipulation of the cells on the day of imaging. 1*µ*L of 2mM VF2.1.Cl in DMSO was mixed with 1*µ*L of 10% Pluronic F127 (diluted in water) in a microcentrifuge tube by agitating and centrifuging 3 times. Mixing of VF2.1.Cl with Pluronic F127 prior to subsequent dilution in physiological buffer is critical for VF2.1.Cl loading. Separately, Hoechst 33258 was diluted into Tyrode’s solution (136mM NaCl, 5mM KCl, 2mM CaCl2, 1mM MgCl2, 10mM glucose, 10mM HEPES, pH 7.4) to a concentration of 4*µ*g/mL. 1 mL of the Hoechst/Tyrode’s solution was added to the the microcentrifuge tube containing the VF2.1.Cl/Pluronic F127 mixture and vortexed for 10 seconds. The contents of the a microcentrifuge tube was added to 4 mL of Hoechst/Tyrode’s solution. Isoproterenol (Sigma-Aldrich) was diluted in DMSO. Each compound was diluted in Tyrode’s solution to a 2x concentrated stock and was warmed to 37°C using a dry heat block prior to application to the cells. (Compounds isoproterenol (Sigma cat# -I6504), Isradipine (Tocris cat#2004), tetrodotoxin citrate (Tocris cat#1069), and dofetilide (Selleck Cat#S1658) were diluted in DMSO. Each compound was diluted in Tyrode’s solution to a 2x concentrated stock and was warmed to 37°C using a dry heat block prior to application to the cells.)

### 2.13. Loading of VF2.1.Cl dye solution and automated image acquisition

On day 30 of differentiation CMs were dissociated with TrypLE Select (Life Technologies) and plated onto Matrigel coated 384-well TC plates (Greiner Bio-One) at a density of 20,000 cells/well. Physiological imaging of cells was performed on day 33 post differentiation.

hiPSC-CMs were removed from a 37°C 5% CO2 incubator and placed immediately onto 37°C warming plate (Fisher) in a tissue culture hood to prevent temperature fluctuation during the subsequent washing and dye loading procedure. Tyrode’s solution was warmed to 37°C prior to adding to a media reservoir located on the 37°C plate warmer. The addition of the VF2.1 dye solution was added following a previously published protocol (McKeithan et al., 2017). After the dye loading period, subsequent washes with Tyrode’s solution to remove any remaining dye and the 2X reference compound was added to each respective well while on the 37°C plate warmer and then the plate was placed back into a 37°C 5% CO2 incubator for 5 minutes to allow for temperature equilibration. The plate was then transferred to a 37°C temperature controlled high content imager (Image Xpress Micro XLS platform, Molecular Devices) and allowed to equilibrate for 10 minutes prior to image acquisition. A single Hoechst image was acquired to autofocus the imager prior to image acquisition. Time series images were acquired using the Image Xpress Micro XLS at an acquisition frequency of 100 Hz for a duration of 10 seconds with excitation wavelength of 485/20 nm and emission filter 525/30 nm using a 0.75 NA Nikon objective. Data was processed using MetaXpress and the physiological parameters were determined.

### 2.14. Statistical analyses

All results shown as averages of triplicates presented as mean ± SEM unless otherwise noted. P values were calculated using a Student’s two-tailed *t* test; *, P value between 0.01 and 0.05; **, P value between 0.001 and 0.01; ***, P value <0.001. All errors associated with ISA were 95% confidence intervals determined via confidence interval analysis. *, statistically significance indicated as non-overlapping confidence.

## 3. Results

### 3.1. Cardiac differentiation increases glucose oxidation of hPSCs

We used an established protocol to differentiate H9 and IMR90iPSC hPSCs into cardiac troponin T-positive (cTNT+) CMs (Burridge et al., 2014). hPSC-CMs were differentiated and maintained in chemically defined CDM3 medium (RPMI1640 containing 500 μg/ml *O. sativa*– derived recombinant human albumin (rHA) and 213 μg/ml L-ascorbic acid 2-phosphate) and either CHIR99021 or WntC59 to modulate Wnt/β-catenin signaling to promote cardiac lineage specific differentiation. To obtain high purity CM cultures, glucose was replaced with lactate for one week to select for cardiac cells (Tohyama et al., 2013). We achieved high efficiency CM differentiation, with over 80% cTNT+ cells after 21 days of differentiation (Fig 1A), which increased further upon lactate selection (Fig 1B). As proper metabolic function is critical for *in vitro* development of hPSC-derived CMs, we subsequently investigated the metabolic features of CMs continually cultured in serum free CDM3 media.

**Figure 1.**
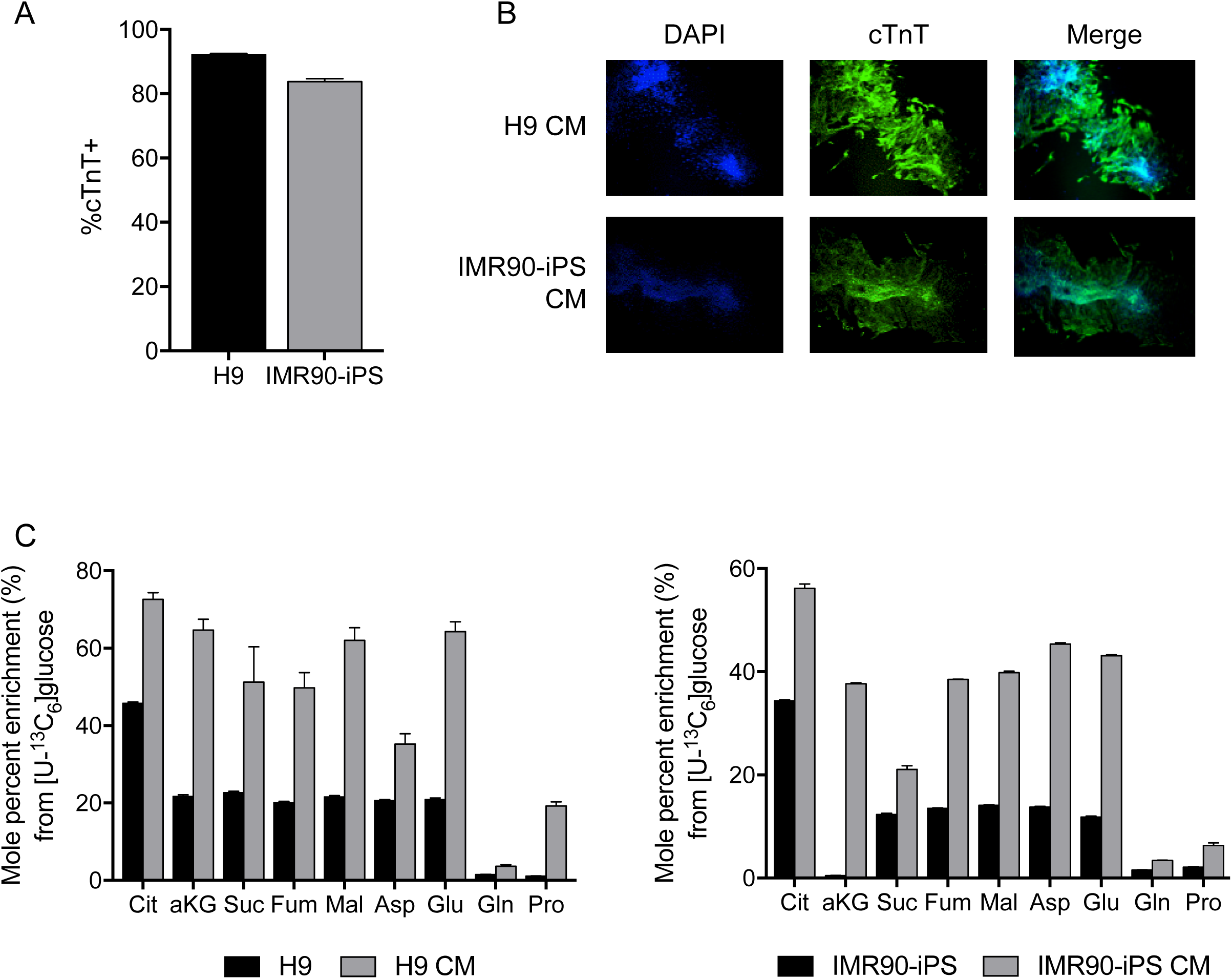
hPSC-derived cardiomyocytes primarily oxidize glucose. (A) cTNT+ flow cytometry of D21 differentiating CMs. (B) Immunofluorescent images of lactate-selected CMs. (C) Central carbon metabolite enrichment from [U-^13^C_6_]glucose in undifferentiated and differentiated H9 (left) and IMR90-iPS (right) cells.

We recently applied ^13^C/^2^H metabolic tracing to demonstrate that hPSCs primarily fuel TCA metabolism using glutamine rather than glucose, as the latter is shunted toward lipid biosynthetic pathways under most hPSC culture conditions (Zhang et al., 2016). Here we similarly quantified how [^13^C]glucose was enriched in TCA intermediates and associated amino acids in hPSCs versus hPSC-CMs. Upon terminal differentiation to the cardiac lineage, we observed a significant increase in glucose oxidation within mitochondria (Fig 1C). Therefore, while glutamine-mediated anaplerosis is important for maintaining hPSCs in the undifferentiated state, differentiated CMs exhibit increased mitochondrial glucose metabolism. However, the dependence on glucose oxidation to generate citrate (above 50%) also suggested that hPSC-CMs differentiated using this approach are metabolically immature, since human adult CMs only exhibit limited glucose contribution to TCA substrates (Lopaschuk and Spafford, 1990, Lopaschuk et al., 1991).

### 3.2. Mitochondrial substrate oxidation in hPSC-CMs

Cardiac tissue is metabolically active and requires efficient nutrient consumption to meet the significant bioenergetic demands of beating CMs (with full ATP turnover occurring over 6 times per minute) (Jacobus, 1985). Mature cardiac cells produce energy from multiple substrates, including fatty acids, glucose, lactate, pyruvate, ketone bodies and BCAAs (Li et al., 2017, Bartelds et al., 1998). As such, hPSC-CMs should exhibit activation of these specific pathways upon differentiation. Importantly, the nutritional microenvironment *in vivo* is vastly different than that of *in vitro* culture conditions used for hPSC-derived CM differentiation, which could limit the metabolic maturation of these cells (Cantor et al., 2017).

We therefore evaluated whether these derivatives would efficiently consume substrates that commonly fuel cardiac metabolism by tracing individual cultures with uniformly labeled [^13^C]glucose, [^13^C]pyruvate, [^13^C]lactate, [^13^C]glutamine, [^13^C]leucine, [^13^C]β-hydroxybutyrate and [^13^C]palmitate tracers. We formulated CDM3 media with these nutrients to measure [^13^C] enrichment in citrate and observed significant incorporation from most nutrients (Fig 2). Notably, β-hydroxybutyrate was significantly oxidized and supplanted a significant quantify of glucose flux into the TCA cycle. On the other hand, the efficiency of fatty acid oxidation was relatively low, as [^13^C]palmitate contributed minimally to citrate in these cultures (Fig 2). Since lipid oxidation is a feature of mature CMs (Khairallah et al., 2004), our results suggested that hPSC-CMs may induce expression of cardiac markers and metabolic enzymes but not effectively activate flux through specific metabolic pathways.

**Figure 2.**
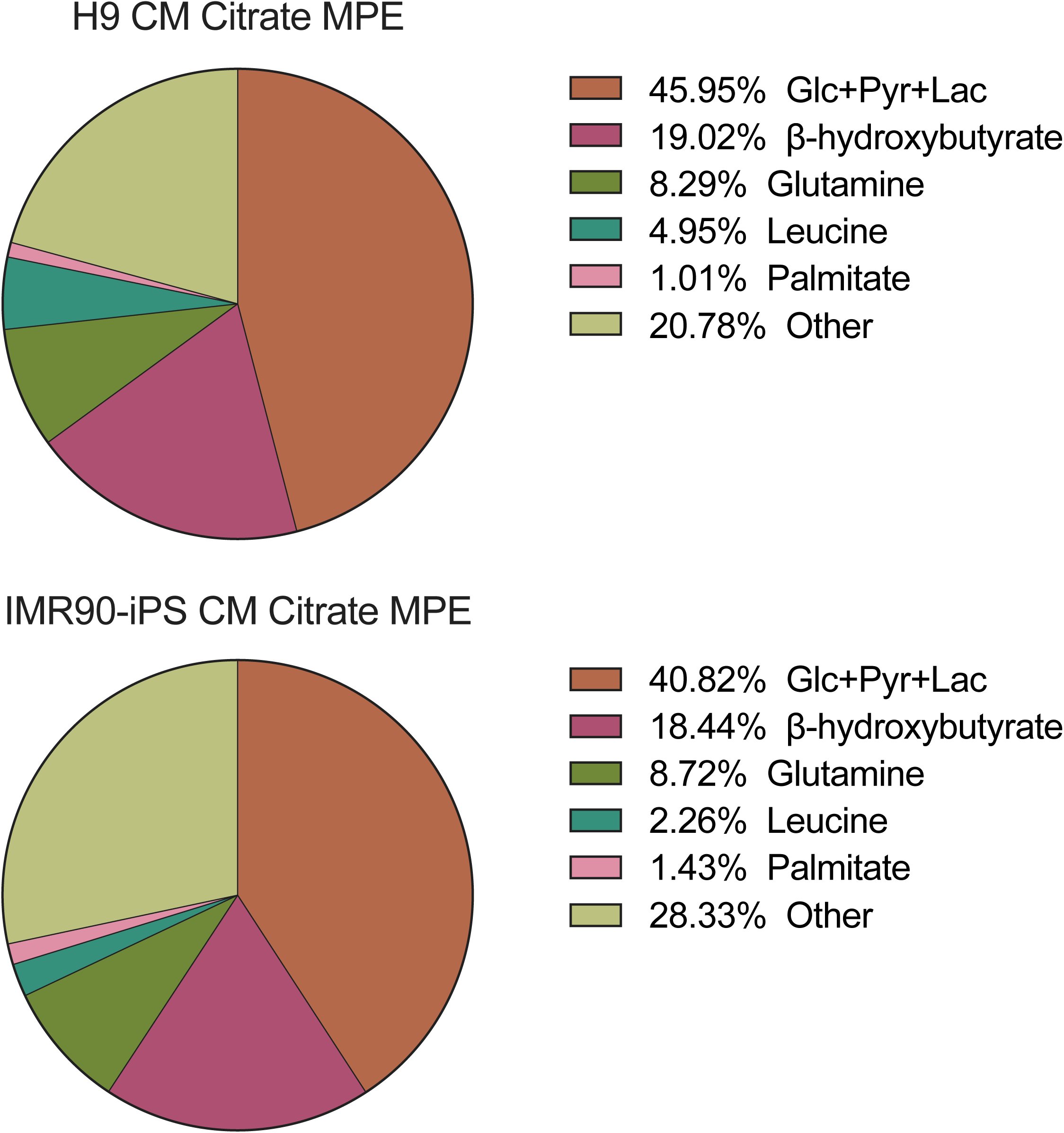
hPSC-derived cardiomyocytes are metabolically immature. (A) Central carbon metabolite enrichment from [U-^13^C_6_]glucose and [U-^13^C_5_]glutamine in H9-derived (left) and IMR90-iPS-derived (right) CMs. (B) Citrate mole percent enrichment from various ^13^C tracers in complex media; H9-derived (top) and IMR90-iPS-derived (bottom) CMs.

### 3.3. Metabolic reprogramming during hPSC cardiac differentiation

To further investigate how metabolic pathway flux and enzyme expression changed during hPSC-CM differentiation we quantified substrate contributions to citrate and pathway-specific gene expression during the first 12 days of hPSC cardiac differentiation. Specifically, cells were maintained in CDM3 media and traced with designated ^13^C-labeled substrates for 24 hours prior to collection of samples for transcriptional and metabolomics analyses at designed time points (Fig 3). As expected in our more replete culture condition, we observed a significant decrease of glucose and glutamine consumption over time (Fig 3A, B). On the other hand, differentiating hPSCs significantly increased oxidation of leucine and β-hydroxybutyrate throughout the cardiac differentiation program (Fig 3C, D). Consistent with the observed changes in substrate oxidation to citrate, we observed that LDH enzyme expression decreased, while expression of the oxidative pentose phosphate pathway enzyme *G6PD* was slightly increased (Fig 3E). Although the expression of both LDH isozymes was slightly decreased, *LDHA* expression decreased more than the heart-specific isoform *LDHB* (Fig 3E). Glutamine contribution to TCA metabolism decreased markedly (Fig 3B), with *GLS2* expression decreasing significantly and *GLS* expression increasing (Fig 3F). Importantly, *GLS2* expression is specifically upregulated in hPSCs maintained in chemically defined media and is presumably more important for hPSC growth and expansion (Zhang et al., 2016).

**Figure 3.**
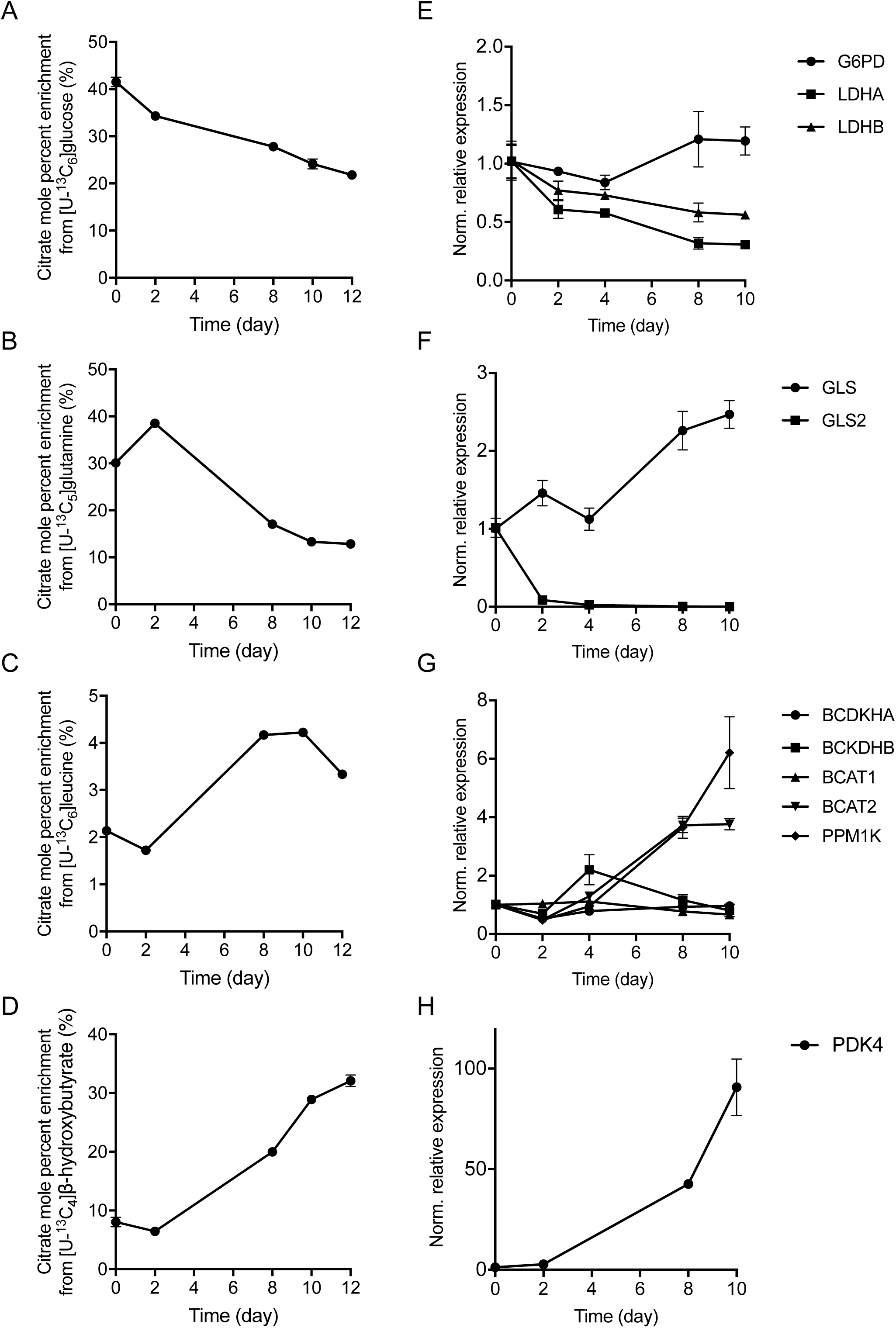
Day-by-day tracing reveals metabolic pathway activation and suppression during cardiac differentiation. Citrate mole percent enrichment from (A) [U-^13^C_6_]glucose, (B) [U-^13^C_5_]glutamine, (C) [U-^13^C_6_]leucine, and (D) [U-^13^C_4_]β-hydroxybutryrate during cardiac differentiation. Tracer added at specified day and metabolites extracted after 24 hours. (E-H) Metabolic gene expression during cardiac differentiation.

In contrast to glutamine metabolism, BCAA oxidation increased significantly, as demonstrated by the leucine contribution to citrate (Fig 3C). Expression of enzymes involved in BCAA catabolism were upregulated with increases in mitochondrial isoform *BCAT2* but not cytosolic isoform *BCAT1* observed (Fig 3G). In addition, the mitochondrial phosphatase *PPM1K*, which promotes branched chain keto acid dehydrogenase complex (BCKDHC) activity through dephosphorylation, was also significantly upregulated (Fig 3G). Furthermore, the significant increase in β-hydroxybutyrate oxidation indicated that ketone body metabolism was highly upregulated during cardiogenesis (Fig 3D). Comparing to the decrease of glucose oxidation (Fig 3A), these results suggested the lower efficiency of cells to produce acetyl-coenzyme A (AcCoA) via pyruvate dehydrogenase (PDH). Importantly, *PDK4*, which negatively regulates PDHC through phosphorylation, was highly upregulated (Fig 3H). These results indicated that hPSCs undergoing cardiac differentiation exhibited reprogramming of mitochondrial metabolism.

### 3.4. Changes in lipid metabolism during hPSC cardiac differentiation

We previously described high rates of *de novo* lipogenesis sustaining the needs highly proliferating hPSCs cultured in chemically-defined media, and hPSC cardiac differentiation is commonly performed in similarly defined media (Zhang et al., 2016). We therefore hypothesized that lipid metabolism might change significantly during cardiogenesis due to the switch of cell behavior from proliferation to differentiation. We first quantified the extent of fatty acid synthesis during differentiation and observed a dramatic decrease in *de novo* fatty acid synthesis as cells became committed to the cardiac lineage (Fig 4A). Consistent with the citrate enrichment data noted above (Fig 3A), hPSCs decreased the contribution of glucose-derived AcCoA into lipid synthesis during cardiac differentiation (Fig 4B).

**Figure 4.**
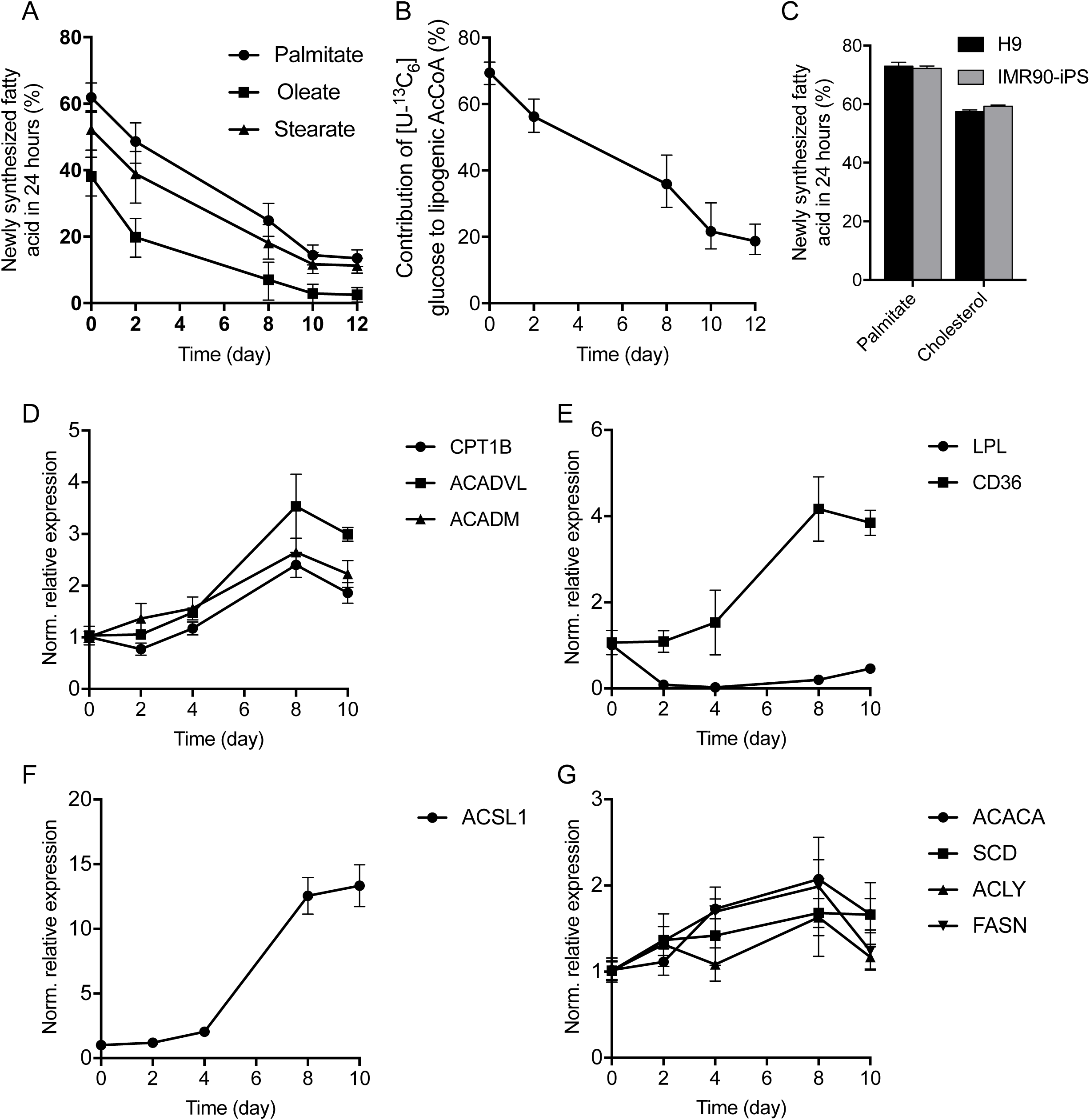
*De novo* lipogenesis is suppressed during cardiac differentiation. (A) Percent newly synthesized fatty acid in 24 hours during cardiac differentiation. (B) Contribution of [U-^13^C_6_]glucose to lipogenic AcCoA during cardiac differentiation. (C) Percent newly synthesized palmitate and cholesterol in 24 hours in hPSCs. (D-G) Fatty acid synthesis and β-oxidation gene expression during cardiac differentiation.

Since nutrient lipids were largely absent from serum-free media, hPSCs exhibited high rates of *de novo* lipid synthesis (Fig 4C). The lack of available lipids in cardiac differentiation media caused us to next consider how the expression of enzymes and transporters involved in lipid utilization might change. Indeed, the expression *CPT1B* (the rate-limit enzyme of long-chain fatty acid β-oxidation pathway), *ACADVL* (the first step enzyme in β-oxidation), and *ACADM* (enzyme in medium fatty acid oxidation) were all significantly upregulated (Fig 4D). At the same time, expression of *CD36* and *ACSL1*, which plays important role in lipid biosynthesis and degradation, were also both increased (Fig 4E-F). In contrast, genes encoding lipogenic enzymes did not change appreciably, including *FASN, ACLY, ACACA* and *SCD* (Fig 4G).

These results suggested that hPSC-CMs differentiated in lipid-deficient conditions might be unable to adequately synthesize or oxidize lipids necessary for maturation or energy generation (Joshi and Sidbury, 1975). Consequently, the maturation of hPSC-CM might be artificially suppressed in serum-free CDM3 media and improved by supplementation of lipids to culture medium.

### 3.5. Immature metabolic features of hPSC-derived CMs cultured in lipid insufficient environment

To test this hypothesis we supplemented hPSC-CM medium with the lipid mixture AlbuMAX on day 10 of differentiation and subsequently compared the differentiation efficiency and metabolic phenotypes to that of control cultures lacking AlbuMAX. AlbuMAX is a lipid-rich bovine serum albumin compromising of a complex mixture of fatty acids (Garcia-Gonzalo and Izpisua Belmonte, 2008). We quantified the fatty acid concentrations in AlbuMAX and associated albumin mixtures to determine what fatty acids were being supplied to CMs (Table 2). Importantly, lipid supplementation did not dramatically impact CM purity after differentiation (Fig 5A). On the other hand, lipid supplementation did not impact CM channel function, as evidenced by their responses to isoproterenol treatment, including increased beating rates and decreased APD90s (Fig 5B) (Lundy et al., 2013, Pillekamp et al., 2012).

**Table 2.**
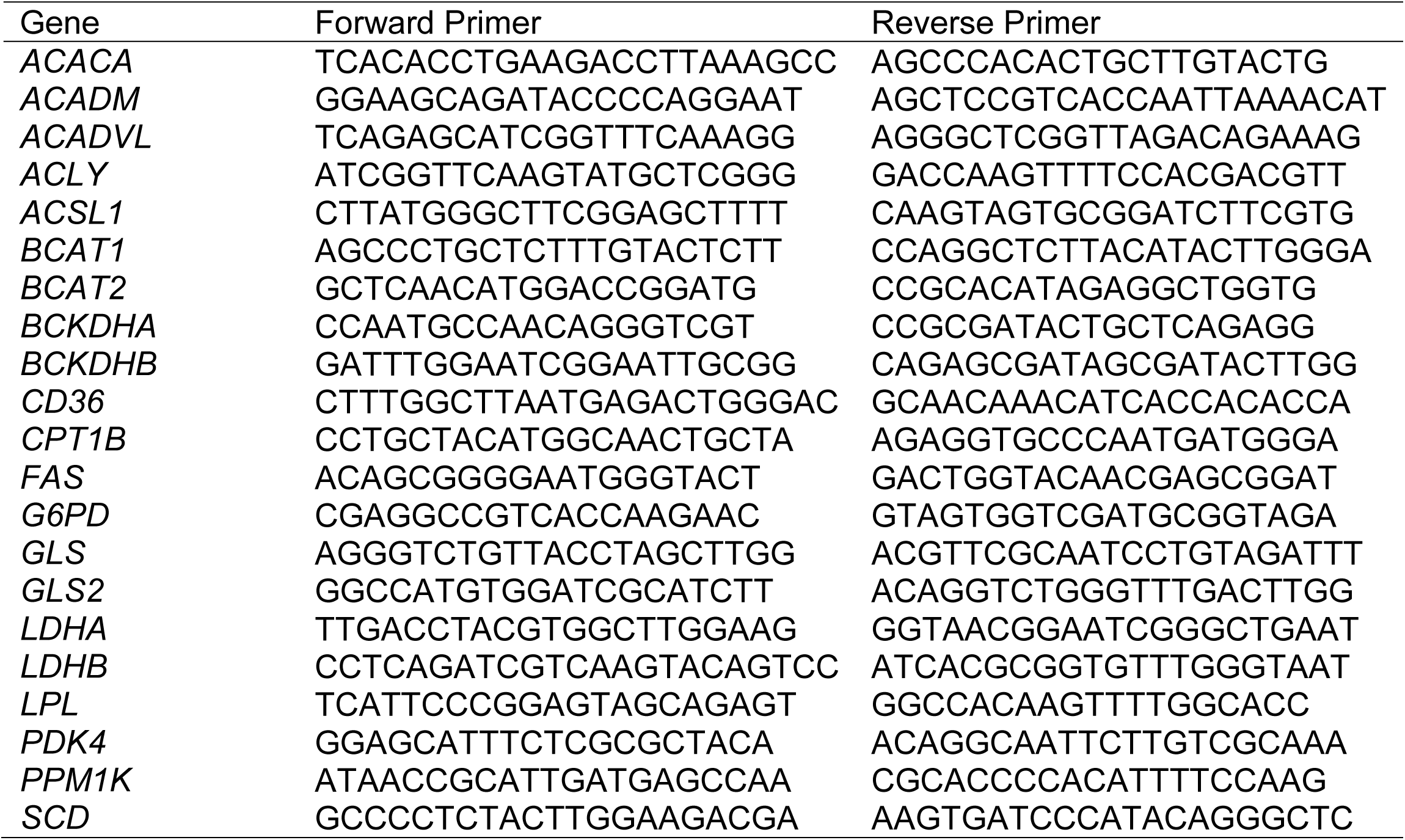
RT-PCR primers.

**Table 3.** Fatty acid concentrations in commonly used albumin media supplements. Data presented as mean ± SD of technical triplicates (pmol/mg albumin).

|  | **Albumax** | **BSA** | **rHA** |
| --- | --- | --- | --- |
| **Myristate (C14:0)** | 570 ± 40 | 1 ± 2 | 0 ± 0 |
| **Pentadecanoate (C15:0)** | 200 ± 30 | 0.65 ± 0.03 | 0.7 ± 0.9 |
| **Palmitate (C16:0)** | 6200 ± 600 | 240 ± 30 | 120 ± 30 |
| **Palmitoleate (C16:1n7)** | 64 ± 8 | 0 ± 0 | 0 ± 0 |
| **Heptadecanoate (C17:0)** | 360 ± 40 | 9.9 ± 0.8 | 0.6 ± 0.7 |
| **Stearate (C18:0)** | 6500 ± 700 | 240 ± 30 | 38 ± 19 |
| **Oleate (C18:1n9)** | 650 ± 80 | 6.1 ± 0.4 | 5.3 ± 1.2 |

**Figure 5.**
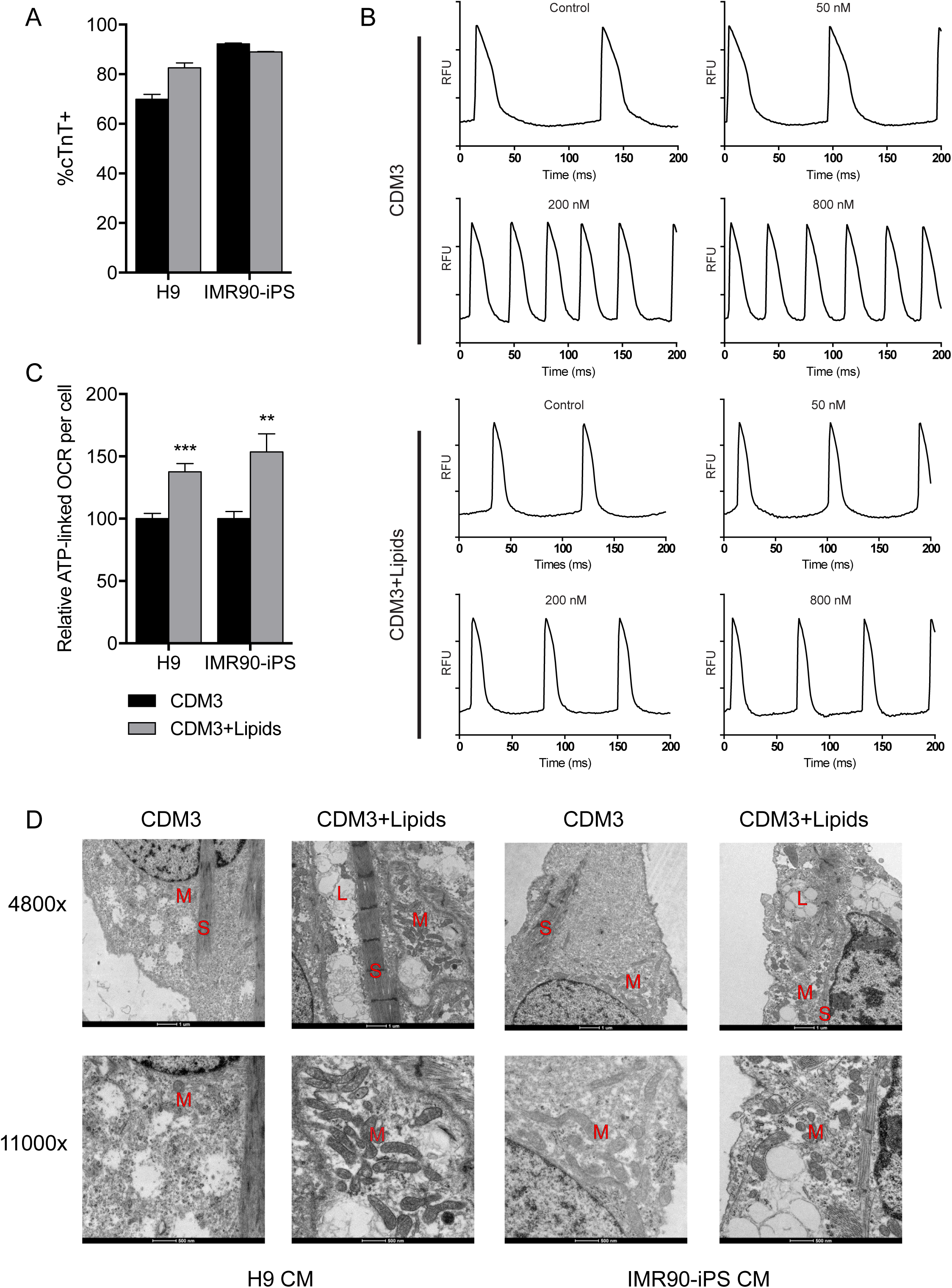
Lipid supplementation activates mitochondrial activity. (A) cTnT+ flow cytometry of CMs cultured with and without lipids. (B) CMs cultured without (top) and with (bottom) lipids increase beating rate in response to beta-adrenergic agonist (Isoprentenol). (C) Relative ATP-linked oxygen consumption for CMs cultured with and without lipids. (D) TEM images of CMs cultured with and without lipids. L – lipid droplet, M – mitochondria, S – sarcomere.

However, we observed a significant increase in ATP-linked respiration in cultures supplemented with AlbuMAX (Fig 5C), indicating that the absence of medium lipids compromised mitochondrial function and oxidative phosphorylation. To further evaluate the impact of lipids on CM cellular development we observed intracellular features using transmission electron microscopy. We observed that hPSC-CMs cultured with lipid supplement present more organized sarcomere structure and larger numbers of mitochondria (Fig 5D). As expected, lipid droplets were only detected in CMs cultured with the AlbuMAX supplement (Fig 5D). As lipid storage, mitochondrial maturation, and sarcomere formation are typical signs of CM development, our results demonstrated that nutrient lipids promote metabolic maturation of hPSC-CMs. These results also indicated that chemically-defined media may not be ideal for the development and long-term maintenance of metabolically mature hPSC-CMs.

### 3.6. Nutrient lipids improve metabolic maturation of hPSC-derived CMs

To further explore the metabolic impacts of lipid supplementation in CM cultures, we examined fatty acid metabolism in hPSC-CMs cultured in the presence or absence of lipid supplements from day 10 to day 28 of differentiation. We first quantified fatty acid abundances of hPSC-derived CMs. Nutrient lipid supplement significantly enhanced cellular fatty acid content (Fig 6A). Odd-chain fatty acids, which are relatively abundant in AlbuMAX as well as milk, increased significantly in supplemented cultures, suggesting these fatty acids were robustly consumed by cells (Fig 6A). The general increase in fatty acid abundance was also consistent with our observation of lipid droplets in supplemented CMs (Fig 5D).

**Figure 6.**
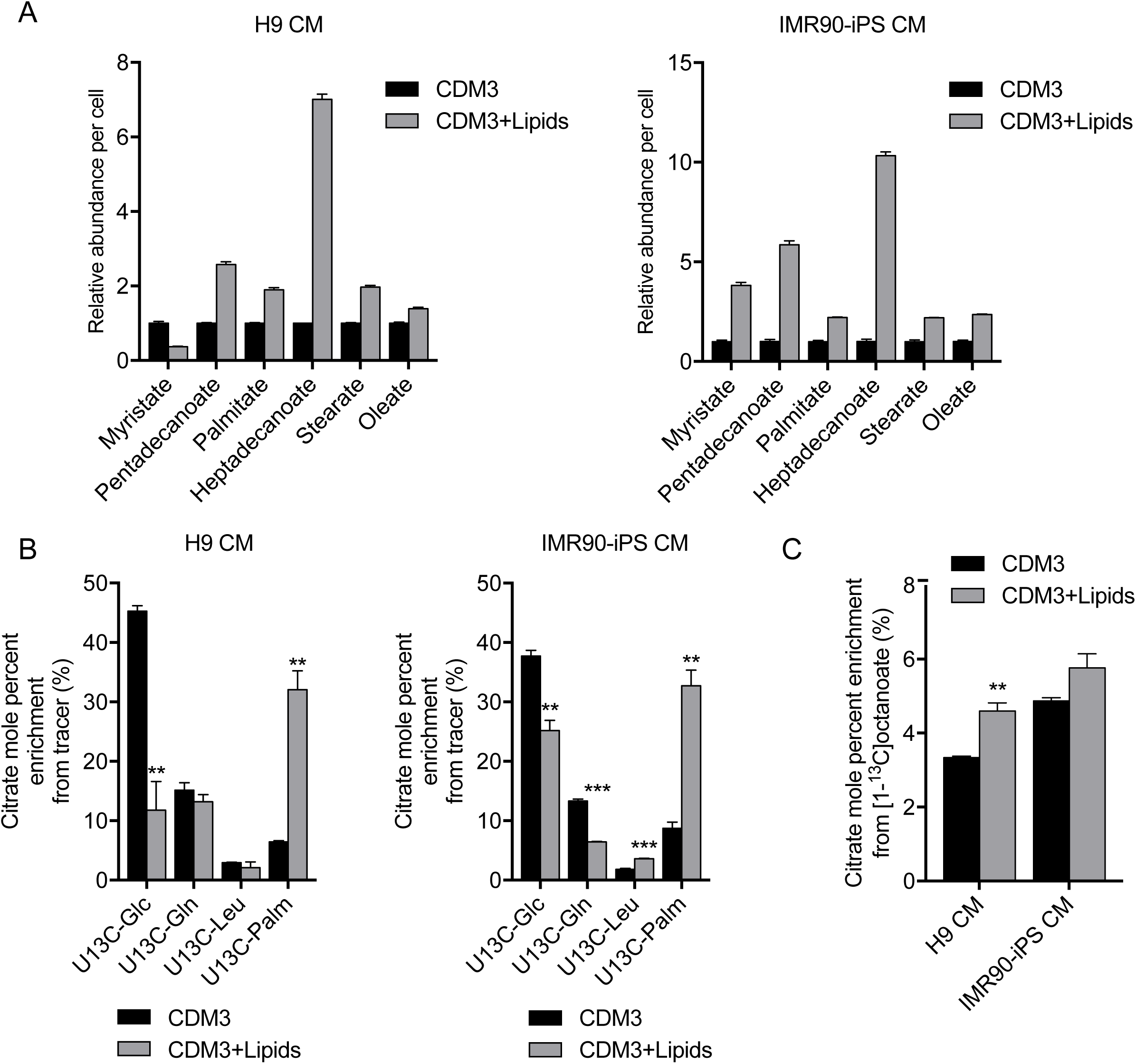
Lipid supplementation increases intracellular fatty acid availability and β-oxidation. (A) Relative intracellular fatty acid abundance per cell in H9 (left) and IMR90-iPS (right) CMs cultured with and without lipids. (B) Citrate mole percent enrichment from specific ^13^C tracer in H9 (left) and IMR90-iPS (right) CMs. β-oxidation of fatty acids is increased when CMs are cultured with lipids. (C) Citrate mole percent enrichment from [1-^13^C]octanoate in CMs cultured with and without lipids.

Next, we examined whether lipid supplementation impacted the extent of fatty acid oxidation in hPSC-CMs. To address this question we cultured hPSC-CMs in the presence of uniformly labeled [^13^C]glucose, [^13^C]glutamine, [^13^C]leucine, or [^13^C]palmitate and quantified isotope enrichment in citrate. We observed that lipid supplementation markedly decreased the contribution of glucose oxidation to TCA metabolism (Fig 6B). Glutamine oxidation was also decreased significantly in IMR90iPSC-CMs (Fig 6B). On the other hand, oxidation of [^13^C]palmitate was significantly increased in AlbuMAX supplemented cultures. To independently confirm this result, we traced cells with [1-^13^C]octanoate, which bypasses the carnitine palmitoyltransferase system, and observed that lipid supplemented hPSC-CMs had increased enrichment in citrate (Fig 6C). These data suggest that nutrient lipids improve the metabolic behavior of hPSC-CMs, presumably through the provision of lipid substrates and biosynthetic intermediates for membrane biogenesis.

## 4. Discussion

Here we have comprehensively profiled the metabolic features of hPSC-CMs during differentiation and maturation. hPSC-CMs are metabolically immature in that they do not effectively oxidize exogenous fatty acids but possess the ability to metabolize other substrates (e.g. ketone bodies and BCAAs). Change in flux correlated with the expression of specific metabolic enzymes and transporters but indicated that cells were deficient in lipids. Exogenous supplementation of fatty acids and lipids improved cardiac morphology, mitochondrial respiration, and FAO while maintaining proper electrical activity.

The pursuit of chemically-defined media for homogenous hPSC maintenance and efficient differentiation has motivated the development of minimal media. While these conditions can support expression of lineage-specific markers and mechanical activity, our data indicates that such culture conditions have metabolic consequences for the cell. Hallmarks of the metabolic maturation of cardiomyocyte include decreases in glycolysis and increases in fatty acid oxidation to regenerate ATP. This process is promoted by mitochondrial biogenesis and activation, which requires structural and signaling lipids (Lopaschuk and Jaswal, 2010). Furthermore, maturing CMs must develop sarcoplasmic reticulum for proper calcium handling, which also requires lipids for membrane biosynthesis. On the other hand, as CMs differentiate and mature, increased *CPT1B* expression requires decreased DNL to prevent malonyl-CoA-mediated suppression of FAO (Saggerson, 2008, van Weeghel et al., 2018). This is exemplified in the heart by the dramatic decrease in ACC and increase in MCD activity during post-natal development (Lopaschuk and Jaswal, 2010, Reszko et al., 2004). Therefore, FAO and DNL are antagonistic processes in the maturing heart and CMs are reliant on exogenous fat sources *in vivo*. Additionally, some chemically defined media (including the CDM3 used here) lack essential ω-3 and ω-6 fatty acids needed for proper lipid composition and cardiac function (Duda et al., 2009, Spector and Yorek, 1985). These essential fatty acids are needed for cardiolipin production, important for proper mitochondrial biogenesis/fusion, and likely promote oxidative metabolism (Paradies et al., 2014), further suggesting that media deficiencies can suppress CM maturation *in vitro* and require further consideration of supplements needed for proper cellular development.

Our approach of providing exogenous nutrients through complex, animal-derived supplements provides one potential avenue to address these issues. Other works have successfully used more diverse nutrient conditions in CM differentiation through defined cocktail addition (Pei et al., 2017, Lian et al., 2012). Cellular physiology must guide the development of differentiation media that promote mature cell performance. Metabolic flux analysis has long been used to identify industrially-relevant cellular bottlenecks (Long et al., 2018) and now is being applied to understand alterations in hPSCs (Zhang et al., 2016, Badur et al., 2015) and cancer (Cantor et al., 2017). Metabolic requirements of non-traditional substrates that support cellular growth is an emerging concept (Pavlova et al., 2018, Sousa et al., 2016) and has already been used to improve cellular differentiation (Beyer et al., 2018). For example immature hPSC-derived CMs showed a preference for oxidation of β-hydroxybutryrate when supplied in our culture conditions, consistent with the physiology of the heart, and should be explored as potential maturation agent (Nagao et al., 2016, Gormsen et al., 2017). Indeed, galactose and fatty acid supplementation has already been utilized to enhance and demonstrates the utility of these approaches (Correia et al., 2017). Our results further demonstrate that environmental nutrient conditions can drive influence maturation of hPSC-derived CMs and promote improved metabolic phenotypes *in vitro*.

## Acknowledgements

The authors acknowledge members of the Metallo lab for technical assistance and helpful discussions. This research was supported by the California Institute of Regenerative Medicine grant (RB5-07356), National Institutes of Health grant R01CA188652, a Searle Scholar Award all to C.M.M., and a NSF Graduate Research fellowship (DGE-1144086) to M.G.B. The authors declare no financial or commercial conflict of interest.

## Author Contributions

H.Z., M.G.B., A.N.M., M.M., and C.M.M. designed the study. H.Z., M.G.B., S.S., A.D., and N.M., carried out the experiments and analyzed data. H.Z., M.G.B., and C.M.M. wrote the manuscript, and all authors read and commented on the paper.

